# Recombination Suppression is Unlikely to Contribute to Speciation in Sympatric *Heliconius* Butterflies

**DOI:** 10.1101/083931

**Authors:** John W. Davey, Sarah L. Barker, Pasi M. Rastas, Ana Pinharanda, Simon H. Martin, Richard Durbin, Richard M. Merrill, Chris D. Jiggins

## Abstract

Mechanisms that suppress recombination are known to help maintain species barriers by preventing the breakup of co-adapted gene combinations. The sympatric butterfly species *H. melpomene* and *H. cydno* are separated by many strong barriers, but the species still hybridise infrequently in the wild, with around 40% of the genome influenced by introgression. We tested the hypothesis that genetic barriers between the species are reinforced by inversions or other mechanisms to reduce between-species recombination rate. We constructed fine-scale recombination maps for Panamanian populations of both species and hybrids to directly measure recombination rate between these species, and generated long sequence reads to detect inversions. We find no evidence for a systematic reduction in recombination rates in F1 hybrids, and also no evidence for inversions longer than 50 kb that might be involved in generating or maintaining species barriers. This suggests that mechanisms leading to global or local reduction in recombination do not play a significant role in the maintenance of species barriers between *H. melpomene* and *H. cydno*.

**Author Summary:** It is now possible to study the process of species formation by sequencing the genomes of multiple closely related species. *Heliconius melpomene* and *Heliconius cydno* are two butterfly species that have diverged over the past 2 million years and have different colour patterns, mate preferences and host plants. However, they still hybridise infrequently in the wild and exchange large parts of their genomes. Typically, when genomes are exchanged, chromosomes are recombined and gene combinations are broken up, preventing species from forming. Theory predicts that gene variants that define species might be linked together because of structural differences in their genomes, such as inverted pieces of chromosomes that will not be broken up when the species hybridise. However, in this paper, we use deep sequencing of large crosses of butterflies to show that there are no long chromosome regions that are not broken up during hybridisation, and no long chromosome inversions anywhere between the two genomes. This suggests that hybridisation is rare enough and mate preference is strong enough that inversions are not necessary to maintain the species barrier.

## Introduction

It is now widely appreciated that the evolution and maintenance of new species is constrained by genetic as well as ecological factors. One major genetic constraint is that, if populations remain in contact, recombination will break down the associations between alleles that characterise emerging species [1]. A large body of work has invoked genetic mechanisms that couple species-specific alleles and so reduce the homogenising effects of gene flow [2, 3]. These include assortative mating, one-allele mechanisms, tight physical linkage, pleiotropy and multiple (or ‘magic’) traits. One mechanism that has been extensively studied is the role of genomic rearrangements such as chromosomal inversions in suppressing recombination in hybrids.

Inversions are widespread in natural populations and are often associated with adaptive traits [4, 5, 6]. Shifts in frequencies of inversions are often associated with ecological niches and clines in natural populations. This has been widely observed across animal taxa including insects (e.g. *Drosophila melanogaster* [7, 8], *D. pseudoobscura* [9, 10] and *Anopheles* mosquitoes [11, 12]) fish (e.g. sticklebacks [13] and cod [14, 15]), birds [16], and mammals [17, 18], as well as in plants (e.g. *Arabidopsis thaliana* [19] and maize [20]) (Table 1). Traits associated with chromosomal inversions include plumage and behavioural traits associated with alternative mating strategies in the ruff *Philomachus pugnax* [21, 22], flowering time and morphological traits in *Mimulus guttatus* [23, 24], colony organisation in fire ants [25], insecticide resistance, host preference and resting behaviour in *Anopheles* [26, 27, 28] and many traits in *Drosophila* [29]. The lack of recombination between inversion haplotypes is thought to maintain multiple co-adapted alleles in complete association, facilitating local adaptation and the maintenance of polymorphism.

**Table 1.** Examples of inversions known to be involved in speciation, adaptation or recombination suppression.

| Species | Inversion | Inversion size<br>(% of chromosome) | Chromosome<br>size | Chromosomes | Genome<br>size | Citation |
| --- | --- | --- | --- | --- | --- | --- |
| *Drosophila pseudoobscura* x *Drosophila persimilis* | Chromosome 2 | 7.6 Mb (25%) | 30.8 Mb | 4 | 171 Mb | [30] |
| *Drosophila pseudoobscura* | Chromosome 3 STPP | 6.6 Mb (34%) | 19.7 Mb | 4 | 171 Mb | [31] |
|  | Chromosome 3 HYSC | 10.7 Mb (54%) |  |  |  |  |
|  | Chromosome 3 STAR | 5.9 Mb (30%) |  |  |  |  |
|  | Chromosome 3 HYST | 8.6 Mb (44%) |  |  |  |  |
|  | Chromosome 3 SCTL | 6.5 Mb (33%) |  |  |  |  |
|  | Chromosome 3 SCCH | 1.7 Mb (9%) |  |  |  |  |
| *Anopheles* species | 2La/2L+<sup>a</sup> | 22 Mb (45%) | 49 Mb | 3 | 278 Mb | [32,33] |
|  | 3La | 22 Mb (54%) | 41 Mb |  |  |  |
| *Arabidopsis thaliana* | Chromosome 4 | 1.17 Mb (6.5%) | 18 Mb | 5 | 135 Mb | [19] |
| *Zea mays* | Chromosome 1 | 50 Mb (16%) | 307 Mb | 10 | 2.1 Gb | [20] |
| *Homo sapiens* | 8p23 | 3.8 Mb (2.6%) | 145.1 Mb | 23 | 3.2 Gb | [34] |
| *Homo sapiens* vs *Pan troglodytes* | 13 pericentric inversions | 165 kb - 78 Mb |  |  |  | [35] |
| *Mus musculus* | t-complex (4 tandem inversions on chromosome 17) | 35 Mb | 95 Mb | 20 | 3.4 Mb | [36] |
| *Gasterosteus aculeatus* (stickleback) | Chromosome I | 442 kb (1.5%) | 28 Mb | 21 | 446 Mb | [13] |
|  | Chromosome XI | 412 kb (2.5%) | 16.7 Mb |  |  |  |
|  | Chromosome XXI | 1.7 Mb (14.5%) | 11.7 Mb |  |  |  |
| *Heliconius numata* | P locus | 400 kb (~4%) | ~10 Mb | 21 | ~300 Mb | [37] |
| *Gadus morhua* (Atlantic cod) | LG1 | 18.5 Mb |  | 23 | 900 Mb | [15] |
|  | LG2 | 5 Mb |  |  |  |  |
|  | LG7 | 9.5 Mb |  |  |  |  |
| *Philomachus pugnax* | Chromosome 11 | 4.4 Mb |  |  |  | [21,22] |
| *Mimulus guttatus* | DIV1 | 6 Mb |  | 14 | 322 Mb | [24] |
| *Solenopsis invicta* |  | 9 Mb |  | 16 | 354 Mb | [25] |
| *Zonotrichia albicollis* | ZAL2 | ~100 Mb |  |  |  | [38] |

Inversions have also frequently been implicated in speciation [39, 40, 41, 42]. Traits associated with reproductive isolation are often linked to inversions [27, 43, 44, 45] and genetic divergence between species can increase within inverted regions through reduction of gene flow [13, 46, 47, 48, 49, 50]. Theory predicts that inversions can spread by reducing recombination between locally adapted alleles [51, 52, 53, 54, 55, 56, 57], which can either establish or reinforce species barriers by capturing loci for isolating traits such as mating preferences and epistatic incompatibilities [58] or allow adaptive cassettes to spread between species via hybridisation [59]. Reduced recombination within and around inversions has been confirmed in several species [35, 47, 60], although it is unlikely that gene flow is entirely suppressed within inversions due to double crossovers and gene conversion [61], factors addressed in some recent models [56, 62].

Several authors have predicted that inversions can enable the formation and maintenance of species barriers in sympatry or parapatry by favouring the accumulation of barrier loci in the presence of gene flow [43, 63, 64, 65]. *Drosophila* species pairs differing by one or more inversions are typically sympatric, whereas homosequential pairs are typically allopatric [43]; in the particular case of *Drosophila pseudoobscura*, sterility factors are associated with inversions in a sympatric species pair but with collinear regions in an allopatric pair [66]. In rodents, sympatric sister species typically have more autosomal karyotypic differences than allopatric sister species [67]. Sympatric sister species of passerine birds are significantly more likely to differ by an inversion than allopatric sister species, with the number of inversion differences best explained by whether the species ranges overlap [68].

However, inversions are not the only mechanism by which recombination rate can be modified during speciation, and more recently attention has been drawn to the potential role of genic recombination modifiers [57]. If recombination rate were systematically or locally reduced in hybrids, this could be favoured in a manner similar to an inversion. Nonetheless, the very strong effect of inversions on recombination rate, and the fact that they are completely linked to the locus at which recombination is reduced, means that they are perhaps the most likely mechanism of recombination rate evolution during speciation.

We set out to test the role of recombination rate evolution in the maintenance of species barriers in *Heliconius* butterflies. The 46 *Heliconius* species allow for a wide range of comparisons at different points along a putative speciation continuum [69, 70, 71, 72]. Chromosomal inversions are known to play an important role in the maintenance of a complex colour pattern polymorphism in *Heliconius numata* [37]. However, no other *Heliconius* inversions are known, due to the lack of resolution in traditional karyotyping [73].

Here, we systematically searched for inversions between sympatric populations of two *Heliconius* species, *H. melpomene rosina* and *H. cydno chioneus*, which are sympatric in the lowland tropical forests of Panama. These species differ by many traits [74] including colour pattern [75], mate preference [76, 77, 78], host plant choice [79], pollen load and microhabitat [80]. Hybrid colour pattern phenotypes are attacked more frequently than parental forms, indicating disruptive selection against hybrids [81]. Assortative mating between the species is strong, and genetic differences in mate preference are linked to different colour pattern loci [78]. Matings between *H. cydno* females and *H. melpomene* males produce sterile female offspring, but male offspring are fertile, and female offspring of backcrosses show a range of sterility phenotypes [82]. Hybrids are extremely rare in the wild, but many natural hybrids have been documented in museum collections [83] and examination of present day genomic sequences indicate that gene flow has been pervasive, affecting around 40% of the genome [84, 85]. Modelling suggests that the species diverged around 1.5 million years ago, with hybridisation rare or absent for one million years, followed by a period of more abundant gene flow in the last half a million years [86, 87], suggesting that the species originated in parapatry, but have been broadly sympatric and hybridising during their recent history.

Models predict that inversions, or other modifiers of recombination, can be established during both sympatric speciation and secondary contact [43, 55, 56, 63, 88]. Furthermore, the genetic basis for species differences between *H. melpomene* and *H. cydno* is well understood and would seem to favour the establishment of inversions. Wing pattern differences are controlled by a few loci of major effect [75], some of which consist of clusters of linked elements. There is also evidence for linkage between genes controlling wing pattern and those underlying assortative mating [78]. The existing evidence for clusters of linked loci of major effect would therefore seem to favour the evolution of mechanisms to reduce recombination between such loci, and hold species differences in tighter association.

We therefore set out to investigate patterns of recombination and test for the presence of inversions between *H. melpomene rosina* and *H. cydno chioneus*. *H. melpomene melpomene* has a high quality genome assembly with 99% of the genome placed on chromosomes and 84% ordered and oriented [89, 90]. Whole genome resequencing has shown that *F*_*ST*_ between *H. melpomene melpomene* and *H. melpomene rosina* is consistently low across the genome, with only a few small, narrow peaks of divergence, but *F*_*ST*_ between *H. melpomene rosina* and *H. cydno chioneus* is substantially higher and heterogeneous [84], and many gene duplications have been identified between the two species [91].

However, *H. melpomene* and *H. cydno* have not yet been examined for evidence of large differences in genome structure. We constructed fine scale linkage maps using multiple crosses of *H. melpomene*, *H. cydno* and *H. cydno* x *H. melpomene* hybrids, to test for the presence of reduced recombination in hybrids and inverted regions between the species (Figure 1). We also generated long read sequencing data and new genome assemblies for both species, to test for inversions on smaller scales. This is the first systematic survey of genome structure and recombination at fine scale in a Lepidopteran species, which may be relevant to the study of holocentric chromosomes and species with no recombination in the heterogametic sex (females in Lepidoptera), in addition to exploring the role of structural variations during speciation.

**Figure 1.**
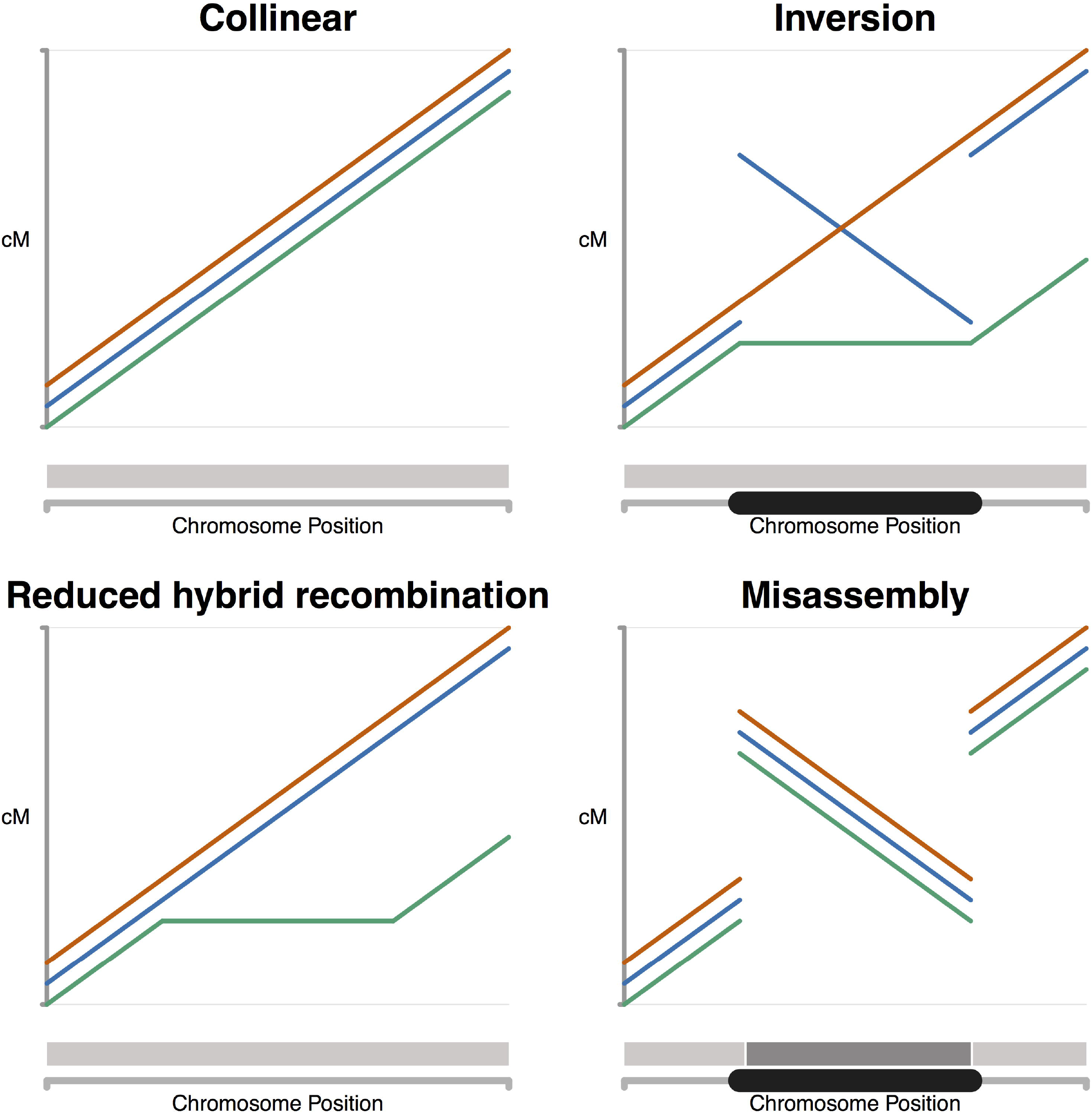
Diagram of expected patterns for collinear, inverted, reduced hybrid recombination, and misassembled genome regions. *H. melpomene*, red; *H. cydno*, blue; *H. cydno* x *H. melpomene* hybrids, green. Grey strip, *H. melpomene* contigs (dark/light grey show different contigs). Black lozenge, inverted region.

## Results

### Summary of crosses and improved ordering of *Heliconius melpomene* assembly

We raised crosses within *H. melpomene*, within *H. cydno* and between *H. cydno* and *H. melpomene*, generated RAD sequencing data for a total of 963 offspring, and built linkage maps for each sequenced cross (Table 2, Table S1, Table S2). We also generated Pacific Biosciences long read data for pools of male and female larvae from *H. melpomene* cross 2 and *H. cydno* cross 1 (Table 3).

**Table 2.**
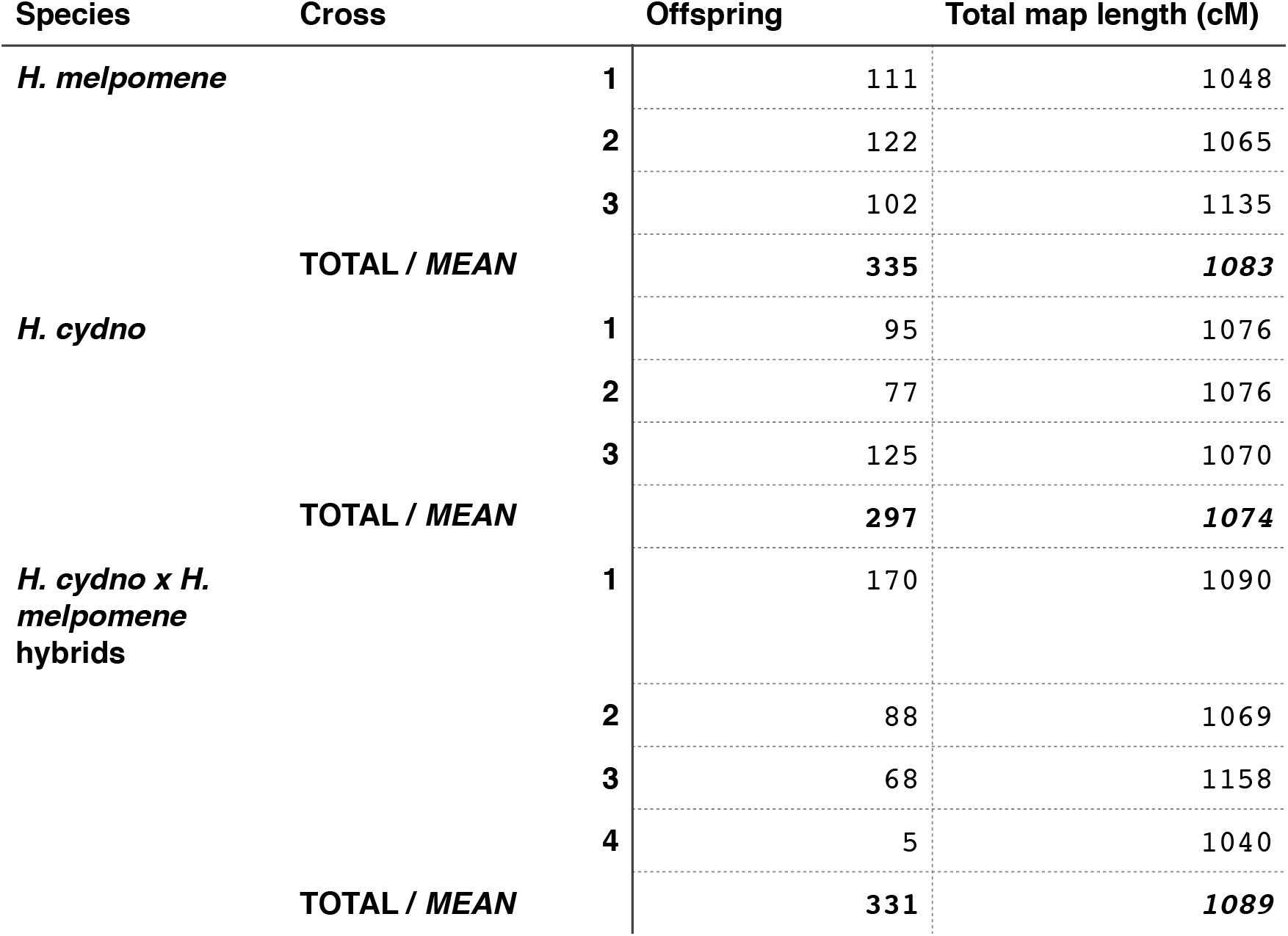
Cross information for each species.

**Table 3.** Summary of Pacific Biosciences sequencing and trio assemblies used to identify inversions.

|  |  | *H. melpomene* |  | *H. cydno* |  |
| --- | --- | --- | --- | --- | --- |
|  |  | Females | Males | Females | Males |
| Pacific Biosciences sequences | SMRT cells | 15 | 10 | 15 | 10 |
|  | Reads | 3,138,554 | 2,079,617 | 3,022,815 | 1,218,186 |
|  | Reads mapped | 3,006,793 | 1,985,294 | 2,745,477 | 1,089,370 |
|  | Reads mapped % | 95.8 | 95.5 | 90.8 | 89.4 |
|  | Bases | 16,172,976,632 | 11,157,098,567 | 15,277,034,979 | 4,457,967,153 |
|  | Bases mapped | 15,842,062,694 | 10,864,609,062 | 14,285,479,974 | 4,170,115,456 |
|  | Bases mapped % | 98 | 97.4 | 93.5 | 93.5 |
|  | Depth mode for mapped bases | 43 | 27 | 38 | 10 |
|  | Bases mapped for PBHoney (primary alignments + tails) | 10,848,364,007 | 7,275,050,641 | 9,173,633,875 | 2,725,704,499 |
|  | Based mapped for PBHoney % | 67 | 65.2 | 60 | 61.1 |
|  | Mode of base depth for PBHoney bases | 37 | 24 | 33 | 8 |
| Trio Assemblies | Scaffolds | 49,035 | 46,134 | 32,548 | 34,566 |
|  | Total Length | 267.8 | 276.8 | 257.9 | 270.3 |
|  | Mean Scaffold Length (kb) | 5.4 | 6.0 | 7.9 | 7.8 |
|  | Scaffold N50 (kb) | 16.9 | 20.1 | 27.0 | 25.7 |
|  | Max Scaffold Length (kb) | 140 | 165 | 234 | 267 |

To improve the accuracy of our recombination rate measurements, we first used the new linkage map of *Heliconius melpomene* and Pacific Biosciences long read data to improve the scaffolding of version 2 of the *Heliconius melpomene* genome assembly (Hmel2; 795 scaffolds, 275.2 Mb total length, 641 scaffolds placed on chromosomes (274.0 Mb), 2.1 Mb scaffold N50 length [90]). This resulted in an updated genome assembly with 13 complete chromosomes, the remaining eight chromosomes having one long central scaffold with short unconnected scaffolds at either end (272.6 Mb placed on 21 chromosomes in 38 scaffolds, including 17 minor scaffolds at chromosome ends totalling 1.3 Mb; 294 additional scaffolds (2.6 Mb) were not placed on chromosomes and unused in further analyses). This updated reference genome assembly was used for all further analyses.

### Differences in recombination rate between species and hybrids

We placed our linkage maps on the new *H. melpomene* chromosomal assembly to look for evidence of reduced recombination in the hybrids. Figure 2 shows Marey maps [92] for each of the 21 *Heliconius melpomene* chromosomes for *H. melpomene*, *H. cydno* and *H. cydno* x *H. melpomene* hybrids, with recombinations from all crosses per species combined (see Figure S1 and Table 2 for per-cross Marey maps and map lengths). Total map lengths are highly consistent between crosses and species across the genome and individual chromosomes (Table 2, Table 4). We designed our crosses with the expectation of finding on the order of one recombination per chromosome per offspring, predicting a recombination on the order of every 45 kb (predicted recombination rate with genetic length 100 cM per chromosome = 8.2 cM/Mb; 292 Mb / 21 chromosomes / 300 offspring = 46.3 kb). However, the actual recombination rate is half of this predicted rate, with a mean genetic length of 51 cM and mean recombination rate of 4.2 cM/Mb per chromosome for both species and hybrids (Table 4) and with no recombinations detected on many individual chromosomes (mean recombinations per offspring across 21 chromosomes in *H. melpomene*, 10.8 (SD 2.4); *H. cydno*, 10.7 (SD 2.2)).

**Figure 2.**
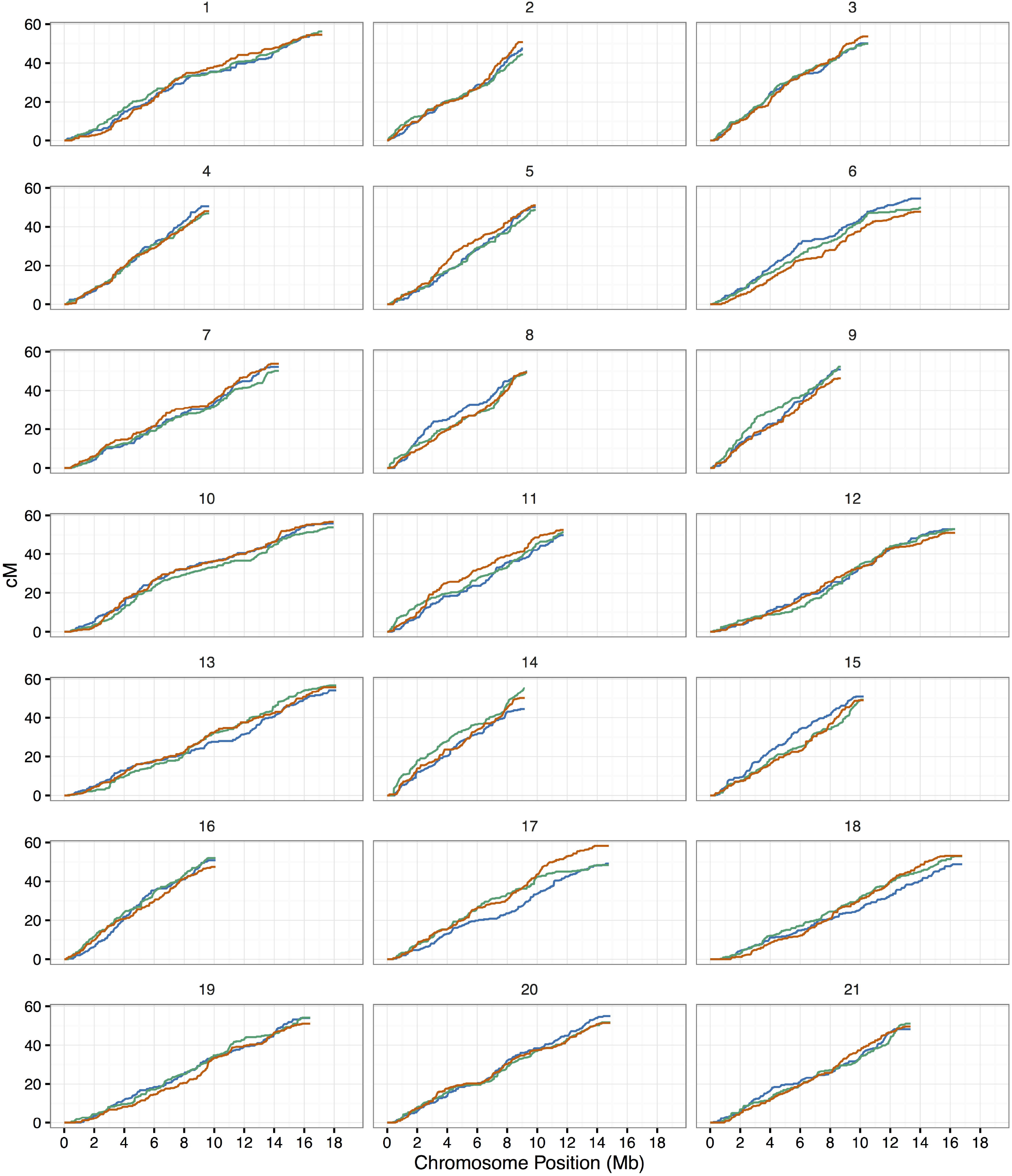
Marey maps of within- and between-species recombination. *H. melpomene*, red; *H. cydno*, blue; *H. cydno* x *H. melpomene* hybrids, green. Chromosomes 1-21 of *H. melpomene* genome assembly version 2 (Hmel2) shown against cumulative centiMorgan (cM) values.

**Table 4.** Physical and genetic map information for each chromosome and species.

| Chromosome | Physical length (bp) | *H. melpomene* |  |  | *H. cydno* |  | *H. cydno x*<br>*H. melpomene* |  |
| --- | --- | --- | --- | --- | --- | --- | --- | --- |
|  |  | Predicted<br>recombination<br>rate (cM/Mb) | Genetic<br>length<br>(cM) | Rate<br>(cM/Mb) | Genetic<br>length<br>(cM) | Rate<br>(cM/Mb) | Genetic<br>length<br>(cM) | Rate<br>(cM/Mb) |
| **1** | 17,206,585 | 5.8 | 54.6 | 3.17 | 54.5 | 3.17 | 56.2 | 3.27 |
| **2** | 9,045,316 | 11.1 | 50.7 | 5.61 | 47.5 | 5.25 | 44.4 | 4.91 |
| **3** | 10,541,528 | 9.5 | 53.7 | 5.10 | 50.2 | 4.76 | 49.9 | 4.73 |
| **4** | 9,662,098 | 10.3 | 48.1 | 4.97 | 50.5 | 5.23 | 46.9 | 4.85 |
| **5** | 9,908,586 | 10.1 | 51.0 | 5.15 | 50.2 | 5.06 | 48.7 | 4.91 |
| **6** | 14,054,175 | 7.1 | 47.8 | 3.40 | 54.5 | 3.88 | 49.9 | 3.55 |
| **7** | 14,308,859 | 7.0 | 53.7 | 3.76 | 52.2 | 3.65 | 50.2 | 3.51 |
| **8** | 9,320,449 | 10.7 | 49.3 | 5.28 | 49.8 | 5.35 | 49.6 | 5.32 |
| **9** | 8,708,747 | 11.5 | 46.3 | 5.31 | 50.8 | 5.84 | 52.3 | 6.00 |
| **10** | 17,965,481 | 5.6 | 56.7 | 3.16 | 55.9 | 3.11 | 53.8 | 3.00 |
| **11** | 11,759,272 | 8.5 | 52.5 | 4.47 | 49.8 | 4.24 | 51.4 | 4.37 |
| **12** | 16,327,298 | 6.1 | 51.0 | 3.13 | 52.9 | 3.24 | 52.9 | 3.24 |
| **13** | 18,127,314 | 5.5 | 55.8 | 3.08 | 54.2 | 2.99 | 56.8 | 3.13 |
| **14** | 9,174,305 | 10.9 | 50.2 | 5.47 | 44.4 | 4.84 | 55.3 | 6.03 |
| **15** | 10,235,750 | 9.8 | 49.0 | 4.78 | 50.8 | 4.97 | 49.3 | 4.81 |
| **16** | 10,083,215 | 9.9 | 47.5 | 4.71 | 50.8 | 5.04 | 52.0 | 5.16 |
| **17** | 14,773,299 | 6.8 | 58.2 | 3.94 | 49.2 | 3.33 | 48.3 | 3.27 |
| **18** | 16,803,890 | 6.0 | 53.1 | 3.16 | 48.8 | 2.91 | 52.9 | 3.15 |
| **19** | 16,399,344 | 6.1 | 51.0 | 3.11 | 53.9 | 3.29 | 54.1 | 3.30 |
| **20** | 14,871,695 | 6.7 | 51.3 | 3.45 | 54.9 | 3.69 | 51.7 | 3.48 |
| **21** | 13,359,691 | 7.5 | 49.6 | 3.71 | 48.1 | 3.60 | 51.1 | 3.82 |
| **GENOME** | **272,636,897** | **7.7** | **1081.2** | **3.97** | **1074.1** | **3.94** | **1077.5** | **3.95** |
| **CHROMOSOME** |  | **8.2** | **51.5** | **4.2** | **51.1** | **4.2** | **51.3** | **4.2** |

Some differences in recombination rate between the species maps are visible; for example, on chromosome 17, the *H. melpomene* map is 9.1 cM longer than *H. cydno*; on chromosome 6, *H. cydno* is 6.8 cM longer than *H. melpomene* (Figure 2, Table 4). However, we are primarily interested in recombination suppression in hybrids, and only chromosome 2 has a significantly reduced recombination rate in the hybrid crosses, and only when compared to *H. melpomene*, not *H. cydno* (one-tailed Kolmogorov-Smirnov tests, see Methods). Regions with reduced recombination in the hybrids can be observed (Figure S2; for example, chromosome 17, 11-13 Mb and chromosome 19, 13.5-14 Mb) but only one region on chromosome 21 shows a significant difference in recombination rate between the hybrids and *H. cydno* (12.3 - 12.8 Mb), with no significant difference between the hybrids and *H. melpomene* and with the hybrid recombination rate increasing, rather than decreasing, in this region (permutation test, see Methods).

### Recombination maps show no major inversions between species

We also examined our recombination maps for evidence of inversions between species (Figure 1). However, there are no regions of any map with a detectable reversed region in *H. cydno* or the hybrids with respect to *H. melpomene* (Figure 2). This is true for the species maps and for all individual cross maps (Figure S1). This indicates there are no large fixed inversions between *H. melpomene* and *H. cydno*.

Known or predicted chromosome inversions involved in the maintenance of species barriers are typically megabases long (Table 1), and models indicate that inversions may have to be very large to become fixed in a population [56]. Our maps are sufficiently fine-scale to rule out the presence of inversions on the megabase scale (*H. melpomene* mean gap between markers, 115 kb, median 87 kb, maximum 1.38 Mb; *H. cydno* mean 135 kb, median 101 kb, maximum 1.14 Mb; see Figure S3, Figure S4). Simulation of random inversions indicates that our existing maps give us power to detect ∼98% of 500 kb inversions, ∼90% of 250 kb inversions and ∼75% of 100 kb inversions (Figure S5). These sizes are smaller than most inversions known to be associated with adaptive traits or species barriers; however, they are on the order of the sizes of the known inversion in *Heliconius numata* and others in, for example, sticklebacks (Table 1). Therefore, we cannot rule out the presence of an inversion in any remaining gap between markers within *H. melpomene* or *H. cydno* from the recombination maps alone.

### Detection of small inversions with long sequence reads and trio assemblies

To test for the presence of smaller fixed inversions between *H. melpomene* and *H. cydno* that were undetectable using our recombination maps, we generated Pacific Biosciences long read sequence data for pools of male and female larvae from one each of the *H. melpomene* and *H. cydno* crosses used to generate recombination maps (Table 3, Figure S6, Figure S7). We called candidate inversions from the long read data using PBHoney to identify reads with clipped alignments, realign the clipped read ends, and detect such split reads with inverted alignments. We also generated Illumina short read assemblies of the maternal and paternal genomes of one offspring from the same crosses used to generate the linkage maps and PBHoney candidates.

These assemblies were constructed using a trio assembly method that separates maternal and paternal reads from one offspring and attempts to construct haplotypic assemblies of each parental genome, providing longer and more accurate contigs compared to standard Illumina assemblies of heterogenous genomes such as those of *Heliconius* species ([93]; Table 3). We aligned these trio assemblies to the *H. melpomene* genome and assessed whether the resulting alignments supported or conflicted with candidate split read inversions.

In total, 1494 raw PBHoney split read candidates were identified across the four samples (two sexes for each of two species; Table 3, Table S3), but many of these were rejected as being incompatible with the linkage maps and trio assemblies. 438 candidates (30%) spanned recombinations in the linkage maps (Table S3), of which 294 (20%) were longer than 1 Mb, with 36 (2.5%) longer than 10 Mb. A further 344 candidates (23%) were removed because the candidate was spanned by a trio scaffold from the same species by 50% of the inversion length on either side (Table S3). These rejected candidates may be false positives; when we simulated PacBio reads directly from the ordered Hmel2 reference genome, PBHoney called 49 ‘false-positive’ inversions. Alternatively, they may be generated by polymorphic inversions, but as we are only interested in fixed inversions, we have not considered these candidates any further.

A further 199 of the 1494 candidates (13%) were removed because they were shorter than 1 kb (Table S3); while these may be real, they are very unlikely to be necessary for species barriers as there is already above-background linkage disequilibrium between SNPs separated by 1 kb or less in *H. melpomene* [94]. The remaining 463 split read candidate inversions from the four samples were merged into 185 candidate groups based on their overlaps. The four samples were sequenced with variable coverage, with low coverage for the *H. cydno* males in particular (Table 3). Given this, 173 additional candidates with less robust support that overlapped with the 185 merged groups were included in the data set (Table S3). Each of the merged groups was then classified based on their presence in either or both species and their support by split read and trio assembly evidence (Table 5, Figure 3, Figures S8-14; see Methods for full criteria).

**Figure 3.**
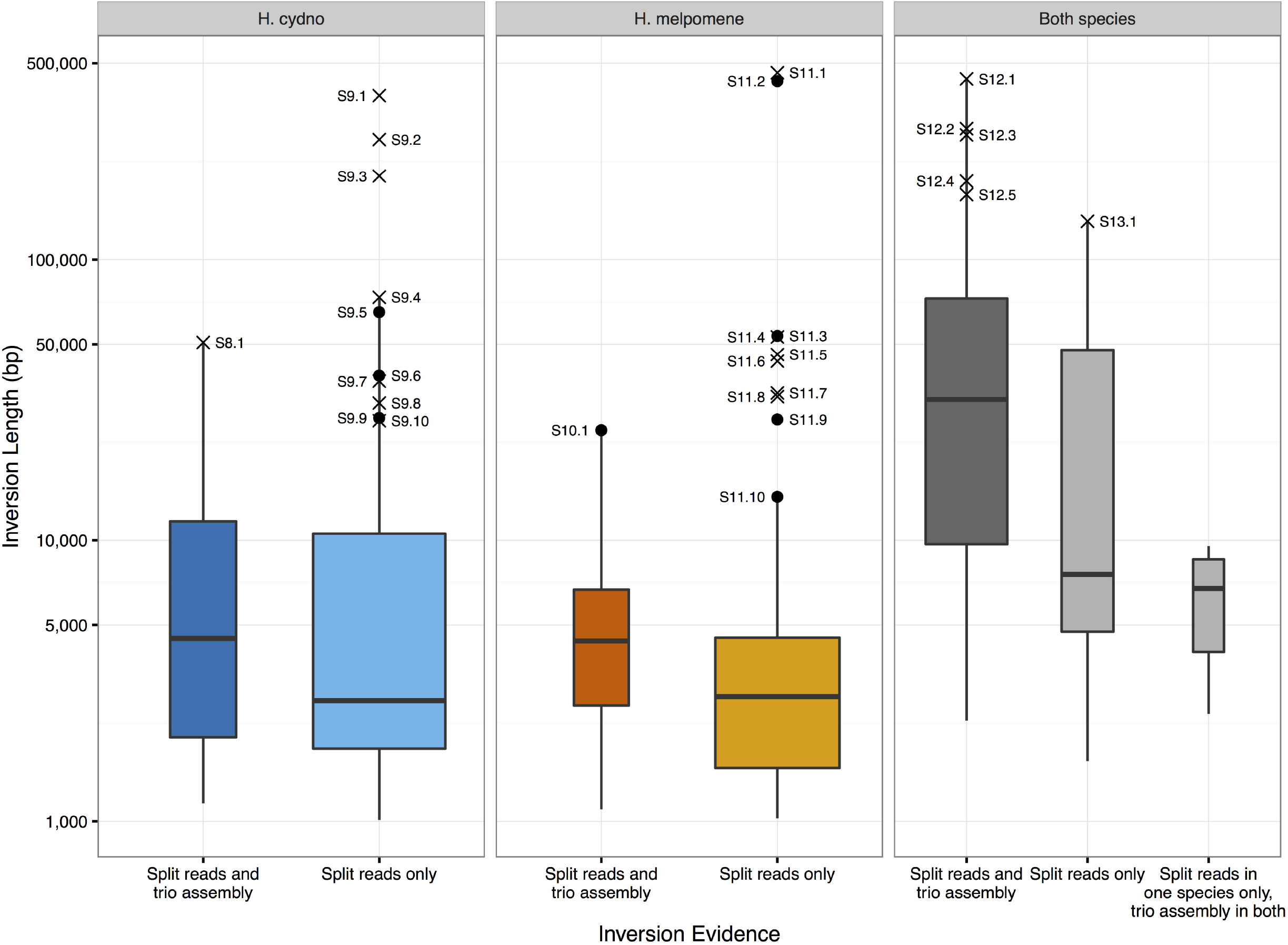
Lengths of candidate inversion groups classified by species and status. See Methods for status definitions. *H. cydno*, blue; *H. melpomene*, red; both species, grey. Dark boxes, evidence from both split reads and trio assemblies; lighter boxes, evidence from either split reads or trio assemblies. Boxes span first and third quartiles; midline shows mean; width represents number of inversions in each category; whiskers extend to the highest value within 1.5 times of the height of the boxes from the edge of the box. Outlier points shown with crosses if contig gaps fall near inversion breakpoints, circles if not. Labels refer to pages of Figures S8-14 where full details of each inversion are given.

**Table 5.** Classification of candidate inversions.

| Split read candidates | Evidence | Candidate inversions | Breakpoint near contig boundaries (%) | Supplemental Figure |
| --- | --- | --- | --- | --- |
| *H. cydno* | Split reads and trio assembly | 13 | 3 (23%) | S8 |
|  | Split reads only | 52 | 17 (33%) | S9 |
|  | **TOTAL** | **65** | **20 (31%)** |  |
| *H. melpomene* | Split reads and trio assembly | 9 | 4 (44%) | S10 |
|  | Split reads only | 46 | 15 (33%) | S11 |
|  | **TOTAL** | **55** | **19 (35%)** |  |
| **Both species** | Split reads and trio assembly | 42 | 39 (92%) | S12 |
|  | Split reads only | 17 | 11 (64%) | S13 |
|  | Split reads in one species, trio assembly in both | 6 | 3 (50%) | S14 |
|  | **TOTAL** | **65** | **53 (82%)** |  |
| **GRAND TOTAL** |  | **185** | **92 (50%)** |  |

Despite the high rate of likely false positives, PBHoney does appear to detect genuine inverted sequences relative to the reference genome, some of which we infer are likely due to local misassemblies in the reference genome. There were 59 groups where PBHoney found overlapping inversions in both *H. cydno* and *H. melpomene*, 50 (85%) of which span multiple contigs, with most inversion breakpoints falling at or near to the end of a contig (Table 5, Figure 3, Figures S12-14). This indicates either that some contigs are inverted, or that the ends of contigs have inverted regions that need to be reassembled (which perhaps explains the failure to fill the contig gaps during assembly). In contrast, candidate inversions specific to either species were less likely to span multiple contigs (20 of 65 *H. cydno* candidates (31%), and 19 of 56 *H. melpomene* candidates (35%); Figures S8-11). We suggest that while some of these species-specific inversions could be explained by misassemblies and incomplete PacBio coverage across both species, many of them could be genuine inversions.

### Candidate inversions assessed using trio assemblies and population genetics

As the false positive rate for PBHoney is high, we made further use of the trio assemblies to find support for the remaining PBHoney candidate inversion groups (Table 5, Figures S8-14). 13 *H. cydno* and 9 *H. melpomene* groups were also supported by trio scaffolds aligning with inverted hits within inversion breakpoints (Figure 3, “Split reads and trio assembly”; Figures S8, S10). Of these, 8 *H. cydno* and 3 *H. melpomene* candidates did not have inversion breakpoints near contig boundaries. If these inversions are species-specific, as indicated by the PBHoney output, we expect support for the reference genome order in the species that does not possess the inversion candidate. Indeed, 6 of these *H. cydno* and all 3 *H. melpomene* candidates have trio scaffolds of the other species spanning the whole inversion or one of the breakpoints, supporting the inversion as being species-specific (Figures S8.2, S8.4, S8.6, S8.8, S8.10, S8.11; Figures S10.6, S10.8, S10.9). However, the longest of these candidates is S8.2 at 20 247 bp, far shorter than any known inversion relevant for speciation, and only slightly larger than the distance at which linkage disequilibrium in *H. melpomene* reaches background levels (∼10 kb [94]). We conclude that there are a small number of likely species-specific inversions, but that these are too small to play a role in speciation. Notably, none of these candidate inversions were located near loci known or suspected to determine species differences in wing pattern or any other trait with known locations [90].

We also calculated *F*_*ST*_ and *f*_*d*_ [95] across inverted regions (Figures S8-S14) to test for differences in the level of admixture at the inversion relative to surrounding regions. An inversion acting as a species boundary typically produces a signal of elevated *F*_*ST*_ and reduced *f*_*d*_ [3, 33, 34, 38, 96], and an inversion enabling the spread of an adaptive cassette might produce a signal of elevated *f*_*d*_ [59]. However, we see very little evidence for increases in these statistics within the handful of candidate inversions compared to the surrounding regions, with only one *H. cydno* inversion (Figure S8.4, 11 719 bp long) showing a noticeable localised increase in *F*_*ST*_. This region contains no annotated features, although of course this does not rule out some functional importance of this region.

Some candidates with only split read evidence, many in only one sex, are hundreds of kilobases long (outliers labelled in Figure 3, particularly those marked with circles, where breakpoints are not near contig boundaries), which, if real, may be relevant to speciation. However, given the large number of false positives produced by PBHoney, the lack of supporting evidence from trio assemblies, and the lack of *F*_*ST*_ and *f*_*d*_ signals at these candidates, it is unlikely these candidates, even if they are real, are substantial species barriers.

### The *H. melpomene* and *H. erato* genomes are mostly collinear, but do contain inverted regions

We have used the recently completed *H. erato* genome assembly to investigate the incidence of inversions between more divergent genomes in the genus. *H. melpomene* and *H. erato* diverged 10 million years ago [71] (Figure S15), considerably more than the ∼1.5 million years between *H. melpomene* and *H. cydno*. Despite the substantial divergence time, the chromosomes of the two species are collinear throughout at the large scale, with a small number of exceptions. There are many regions of the *H. erato* genome assembly that are inverted relative to the ordered Hmel2 assembly, but they fall within single linkage map markers and so may be due to genome misassemblies. For example *H. erato* scaffolds Herato0201, Herato0202 and Herato0203 on chromosome 2, and the first 300 kb of *H. melpomene* chromosome 3, may be misoriented rather than genuinely inverted.

However, three large inverted and/or translocated regions have multiple linkage map markers in both species, and so are likely to be genuine inversions (Figure S15; chromosome 2, *H. erato* 7-10 Mb, *H. melpomene* 4-7 Mb; chromosome 6, *H. erato* 16-18 Mb, *H. melpomene* 12-13 Mb; chromosome 20, *H. erato* 13-15 Mb, *H. melpomene* 11-12 Mb). The chromosome 2 rearrangement, on current scaffold ordering, appears to be an inversion followed by a translocation (for scaffold Herato0214), but given that scaffolds Herato0212, Herato0213 and Herato0214 are all found at the same marker on the linkage map, it may be that these scaffolds need to be reoriented and reordered, so this region could represent a single inversion.

From these comparisons, it is clear that, while some chromosome rearrangements exist, *Heliconius* species do not feature inversions on the scale of, for example, *Drosophila*, where there are over 300 known inversions in *D. melanogaster* and over 50 on the third chromosome alone for *D. pseudoobscura* and *D. persimilis* [97].

## Discussion

We have systematically tested the hypothesis that reduced recombination rates in hybrids might be favoured during speciation with gene flow [57]. High density linkage maps and high coverage long-read sequence data give us considerable power to both measure recombination rate and detect structural rearrangements, but we find no evidence for differences between *H. melpomene* and *H. cydno* at a scale that is likely to influence the speciation process.

Our data have some limitations that might have prevented us from identifying genuine inversions between or within *H. melpomene* and *H. cydno*. Firstly, we have only sequenced crosses from three or four pairs of parents per species, and so may have missed polymorphic inversions due to sampling of wild individuals. However, any inversion important for speciation is expected to be fixed between the species, so it should have been detected even in small samples. Secondly, the maps contain regions of the genome up to a maximum of 1.3 Mb without recombinations that might conceivably harbour inversions (see Results, Recombination maps show no major inversions between species); further crosses could improve resolution in these areas. Recombination rates were half of our naive expectations of one recombination per chromosome per offspring, with individual offspring commonly having no detectable recombinations on half of the 21 chromosomes. This suggests that lepidopteran males do not require recombination for meiosis to proceed, which is perhaps consistent with the fact that meiosis in lepidopteran females occurs without recombination; alternatively, recombination could occur at the ends of chromosomes and be undetectable with our current genome assemblies.

We have found no evidence for significantly reduced recombination in hybrids. Larger crosses would also give greater resolution to this test, and might detect smaller regions of reduced recombination. Nevertheless, we can decisively rule out the presence of any multi-megabase rearrangements among these samples.

We complemented the linkage maps with PacBio sequencing and trio assemblies, in order to detect candidate inverted regions at a smaller scale. This approach also has challenges and generated a high rate of false positives. One potential source of difficulties is reliance on alignment to the *H. melpomene* reference genome assembly. The existing assembly has 25% transposable element content [98] and is likely missing around 6% of true genome sequence, mostly due to collapsed repeats [90]. Inversion breakpoints are typically repeat-rich, which increases the likelihood that reads or scaffolds will not align correctly, and that the breakpoint regions could be misassembled or absent in the reference genome and in the trio assemblies. This problem may be worse for *H. cydno*, where more divergent sequence may align incorrectly or not align at all, and unique *H. cydno* sequence will not be present in the reference (an additional ∼5% of *H. cydno* sequence did not map to the *H. melpomene* genome compared to *H. melpomene* samples (Table 3)). These issues may explain the high observed rate of false positives in our data.

Nonetheless, many of the candidate inversions found in both species are likely to be real with respect to the reference genome, as they are supported by multiple lines of evidence and fall near contig boundaries. Where these are identified in both species, we hypothesise that they reflect misassemblies relative to the reference assembly. These misassemblies could be due to whole inverted contigs, or to misassembled inverted regions at the ends of contigs, which may be preventing the contigs being joined by spanning reads. These examples of misassemblies demonstrate that our methods are capable of detecting large rearrangements in the sampled reads relative to the genome assembly.

In contrast, candidate species-specific inversions are typically smaller than the candidate misassemblies, and are mostly not supported by multiple lines of evidence. Indeed, we can find no compelling fixed candidate inversions supported by both the split read and trio assembly data sets that also show evidence of an increase in *F*_*ST*_, except for the 11.7 kb inversion shown in Figure S8.4, which is too small to substantially increase linkage across this locus beyond that expected by normal decay of linkage disequilibrium [94]. It is possible that some of the candidates with less robust evidence are genuine, given the limitations described above, but on the existing evidence we cannot robustly identify any inversions that are likely to be involved in maintaining species barriers between *H. melpomene* and *H. cydno*.

Although existing models identify situations where chromosome inversions can spread to fixation between two species and maintain a species barrier, the spread of inversions is by no means inevitable during speciation with gene flow. For example, in the model of Feder, Nosil and Flaxman [56], inversions only fix when the strength of selection on the loci captured by the inversion is considerably lower than migration between the species. Similarly, Dagilis and Kirkpatrick [58], modelling the spread of inversions that capture a mate preference locus and one or more epistatic hybrid viability genes, show that inversions are unlikely to spread where pre- and post-zygotic reproductive isolation is already strong. In a recent review, Ortiz-Barrientos et al. [57] also highlight that during reinforcement, assortative mating and recombination modifiers such as inversions are antagonistic; if strong assortative mating arises first, there is only weak selection for reduced recombination.

We considered *H. melpomene* and *H. cydno* to be good candidates for the spread of inversions because there are linked loci causing reproductive isolation and hybridisation has been ongoing for much of their history. Comparisons between sympatric and allopatric populations of the two species have shown that almost a third of the genome is admixed in sympatry [84]. Thus, hybridisation has been ongoing for a long time, perhaps at a low rate. Nonetheless, perhaps strong selection on species differences and assortative mating do not provide the conditions that would have promoted the spread of inversions. Aposematic warning patterns are strongly selected [99], with F_1_ hybrids twice as likely to be attacked as parental phenotypes [81], and pre-zygotic isolation in the form of mate preference is almost complete [76]. Therefore, inversions may not be necessary for divergent loci to accumulate between the species. Perhaps in this case, the evolution of strong assortative mating was favoured by reinforcement selection before inversions or other mechanisms for reducing recombination arose.

This conclusion is also consistent with the apparently low background rate of fixation of inversions in *Heliconius* genomes. *H. melpomene* and *H. erato*, which last shared a common ancestor over 10 million years ago, have largely collinear genomes, and there is substantial chromosomal synteny across the Nymphalids (Ahola et al. 2014). This contrasts with the frequent occurrence of inversions in *Drosophila* genomes, for example, even within single species. The association of multiple inversions with the wing pattern polymorphism in *H. numata* is all the more remarkable given the low background rate of inversions in these butterflies. In conclusion, we have clearly shown that species barriers can persist during speciation with gene flow without suppression of between-species recombination.

## Materials and Methods

Bespoke scripts, data files and further details of methods can be found in the data repository associated with this manuscript. This repository will be submitted to Dryad on acceptance but is currently available on Dropbox at https://www.dropbox.com/sh/aezq5a35f78o03l/AAALDO0kvB1rYMXH8_1PTiIa?dl=0.

### Crosses

Crosses were prepared in Smithsonian Tropical Research Institute insectaries in Gamboa, Panama. Individuals used to establish stock and for crosses were collected from Gamboa (987.4°, 79842.2° W, elevation 60 m) and the nearby Soberania National Park, República de Panamá. For the within-species crosses of *H. melpomene rosina* and *H. cydno chioneus*, wild males were mated to virgin stock females. Females were placed alone in a cage for laying with access to *Psiguria* flowers, *Lantana camara*, and artificial feeders containing a 20% sugar-water solution with 5% added commercial pollen, changed every other day. All laying females were provided with *Passiflora* plants (*H. melpomene*, *P. menispermifolia*; *H. cydno*, *P. edulis*, *P. vitifolia*, *P. triloba*, *P. quadragulata*) with fresh shootings for laying. Eggs were collected every day, and separated into individual plastic pots to prevent cannibalism. Larvae were reared individually on fresh *Passiflora* leaves until late 3rd or early 4th instar, preserved in 2ml of 20% DMSO and 0.25M EDTA (pH 8.0) and stored at −20°C. Adults were preserved and stored in the same way after dissecting wings. Adult males were preserved after mating; adult females were preserved when egg laying ceased. Interspecific crosses were achieved by placing pupae from a *H. cydno* stock in a cage of wild *H. melpomene* males. Hybrid individuals used to produce linkage maps were then obtained from backcrosses produced by mating F1 males to females from the *H. cydno* stock. As with the infraspecific crosses, eggs were collected daily, and larvae were then raised individually; however, unlike the infraspecific crosses, individuals were raised to adults.

### Dissection and DNA extraction

All dissections were performed with a new sterile scalpel, fresh Parafilm and tweezers washed in 80% ethanol. For adults, the thorax was dissected away from the head and abdomen and cut in half along the median plane. One half of thorax was used for DNA extraction with the remaining tissue returned to storage. Larvae were cut in half along the median plane and gut contents removed, with one half used for extraction and the other returned to storage. Some larvae were too small to be cut along the median plane and were cut along a transverse plane instead, whereas some halves of large larvae were cut into smaller pieces to fit within kit tissue size limits.

DNA for all RAD sequenced samples was extracted with the QIAGEN DNeasy Blood & Tissue kit (69504) following manufacturer’s instructions for animal tissue. DNA for Pacific Biosciences sequenced samples was extracted with the QIAGEN MagAttract HMW DNA kit (67563) following manufacturer’s instructions for fresh or frozen tissue, with overnight lysis of tissue. DNA quantity, purity and size distribution and were measured using a Qubit Fluorometer (Q32857), Nanodrop Spectrophotometer (ND-1000) and Agilent 2100 Bioanalyzer (G2939A).

### RAD Sequencing library preparation

RAD libraries were prepared to contain 96 samples in 6 batches of 16, with each sample ligated to one of 16 P1 adapters and each batch ligated to one of 8 P2 adapters (adapters provided by IDT). 500ng of DNA for each sample was diluted to 35uL and digested with NEB high fidelity PstI enzyme (1.0uL PstI-HF, 5.0uL NEB Cutsmart buffer, 9.0uL dH_2_O), incubating for 60 minutes at 37°C and denaturing for 20 minutes at 80°C. Samples were ligated to P1 adapters (add 4ul of 100 nM P1 adapter, 1uL NEB Buffer 2, 0.6uL 100mM rATP, 1.0uL T4 Quick Ligase and 3.4uL dH2O to 50ul DNA solution, incubate for 60 minutes at 22°C and denature for 10 minutes at 65°C), pooled, sheared using a Diagnode Bioruptor (UCD 200 TM) for 8 minutes (30s on, 30s off) and cleaned up with the Nucleospin kit following manufacturer’s instructions (Machery Nagel 740609.50). Libraries were size selected to 250-700bp using a Sage Science BluePippin system (BLU0001) with 1.5% agarose cassettes (BDF1510). 38uL of the resulting library was blunted with the NEB Quick Blunting kit (NEB E1201L) (adding 5uL 10x Blunting Buffer, 5uL 1mM dNTPs, 2uL Blunting Enzyme and incubating for 30 minutes at 30°C) and purified (mix 0.8:1 Ampure XP beads (Beckman Coulter A63880):DNA, vortex briefly, incubate at room temperature for 5 minutes and place in magnetic rack for 3-5 minutes; remove supernatant and wash in 250uL 80% ethanol twice; dry beads, add 42uL elution buffer and resuspend; return to magnetic rack and remove 41uL of eluate).

Libraries were A-tailed (add 5uL 10x NEB Buffer 2, 1uL 100mM dATP, 3uL exo-Klenow fragment (NEB E6044) and incubate for 30 minutes at 37°C, then purify as before to 40.5uL eluate) and P2 adapters were ligated (add 3uL 10uM P2 adapter, 5uL 10x NEB Buffer 2, 0.5uL 100mM rATP, 2uL T4 Quick Ligase (NEB M2200L) and leave for 60 minutes at room temperature); samples were purified as above and quantified, to a final yield of 5-20 ng/uL. Libraries were amplified by PCR using P1 and P2 adapter primers for 12-16 cycles depending on initial yield, and using P2 primers with different barcodes for each batch of 16 P1-ligated samples. Each batch was amplified in 8 separate reactions, to increase diversity of initial samples and reduce sequencing of PCR duplicates, and pooled. Amplified samples were purified as above and eluted in EB-Tween 0.1%.

### RAD library sequencing and alignment

RAD libraries were sequenced by BGI using the Illumina HiSeq 2500 (hybrids and *H. cydno* samples) and HiSeq 4000 (*H. melpomene* samples) platforms. FASTQ files were demultiplexed using process_radtags in Stacks v1.34 [101] allowing 2 mismatches between barcodes. Sequences were aligned to version 2 of the H. melpomene genome (Hmel2 [90]) using Stampy [102] v1.0.23 with options --substitutionrate=0.01, --gatkcigarworkaround, --baq and --alignquals, and then processed by Picard MarkDuplicates (v1.135, http://broadinstitute.github.io/picard/) and GATK IndelRealigner (v3.4.0 [103]). Genotype posteriors were called with SAMtools mpileup (v1.2 [104]) requiring minimum mapping quality of 10 and minimum base quality of 10. A small number of individuals were removed during linkage map construction due to low coverage or large numbers of erroneous genotype calls in markers (light grey samples in Tables S1 and S2).

### Pacific Biosciences library prep and sequencing

Larvae from H. cydno cross 1 and H. melpomene cross 2 were sexed by examining the segregation of the Z chromosome where linkage maps were available at time of sequencing, or the ratio of average autosomal RAD sequencing read coverage to sex chromosome read coverage. Pools of 12 males and 12 females were constructed, using larvae with the highest DNA yields. 12 individuals were used per pool to achieve sufficient yield for Pacific Biosciences sequencing and to approach even coverage of the four parental genomes (insufficient DNA was available to generate long read sequences from parents directly). Each pool was size selected to fragments greater than 6 kb using the SageELF system. Libraries were prepared following the standard 10 kb protocol (SMRTbell Template Prep Kit 1.0, 100-259-100) but with only two bead clean ups at the end rather than three. Libraries were sequenced on a PacBio RSII machine with P6/C4 reagents for 3-4 hour run times. Subreads were used for analyses.

### Linkage mapping: within-species crosses

Within-species linkage maps for *H. melpomene* and *H. cydno* were built with Lep-MAP2 ([105]; downloaded from https://sourceforge.net/p/lepmap2, commit d91aa7, 18 January 2016) and some modules from Lep-MAP3 (noted below; https://sourceforge.net/projects/lep-map3/). For each cross, sites were filtered with Lep-MAP2 pileupParser2.awk and Lep-MAP2 pileup2posterior.awk, requiring each SNP to be sequenced at least 3 times for 90% of individuals, and requiring each allele to be present in at least 5% of individuals. Missing parental genotypes were called with the Lep-MAP2 ParentCall module with options ZLimit=2.0 and removeNonInformative=1.

Sites were clustered into markers by filtering for segregation distortion with the Lep-MAP3 Filtering module with option dataTolerance=0.01 (distortion by chance 1:100) and then processing by LepMAP3 SeparateIdenticals, setting lodLimit options to 20 for maternal markers, log_10_ 2^(*n*-4) for paternal markers and log_10_ 3^(*n*-4) for intercross markers (*n* = number of individuals in the cross; calculation based on 2 possible genotypes for paternal markers, 3 for intercross markers, and allowing for 4 missing individuals), betweenSameType=1, lod3Mode=2, keepRate=1, numParts=2; markers were then passed through Lep-MAP2 OutputData with option sizeLimit=3 (retaining only those markers found at at least three sites). Filtered, clustered markers were then separated into linkage groups using Lep-MAP2 SeparateChromosomes, setting lodLimit empirically for each cross to produce 21 linkage groups. Markers for each linkage group were then ordered with LepMAP2 OrderMarkers, setting maxDistance=0.1, initRecombination=0.05,0.000001, and learnRecombinationParameters=1,0.

Initial marker orderings were manually reviewed and edited to correct misorderings, including removal of low quality markers causing disorder, and to ensure linkage groups were consistent with Hmel2 chromosome directions, using script clean_Lep-MAP_output.pl. Maps were then improved in two directions by script assign_snps_to_markers.pl: i) using Lep-MAP3 module JoinSingles2, all SNPs were reassigned to the set of cleaned markers, to extend the coverage of the linkage map across Hmel2 scaffolds; ii) where two consecutive markers differed by multiple recombinations, all possible paths between the two markers were generated and checked against JoinSingles2 output, with the most likely path being included in the map.

### Linkage mapping: hybrid crosses

Due to more complex cross structure involving F_2_ populations, smaller cross sizes and lower sequence quality for some crosses, different methods were used to construct linkage maps for the *H. cydno* x *H. melpomene* hybrid crosses, now incorporated into Lep-MAP3 (https://sourceforge.net/projects/lep-map3/). SNPs were required to be sequenced at least 3 times for 80% of individuals, requiring 1200 total reads across all individuals and at least two alleles with minimum read coverage of 60 each. Lep-MAP3 module ParentCallGrandparents was called on posterior data with parameters ZLimit=2 and removeNonInformative=1 to impute missing parental calls. Families with less than 3 offspring were removed. Markers with high segregation distortion (p<0.001, distortion by chance 1:1000) were removed with Lep-MAP3 module Filtering2, with parameter dataTolerance=0.001. Markers were first separated to 21 chromosomes with Lep-MAP3 module SeparateChromosomes2 with recombination rate 3% (parameter theta=0.03) and LOD score limit 35 (lodLimit=35). The 21 largest linkage groups were matched to *H. melpomene* chromosomes and additional markers included using Lep-MAP3 module JoinSingles2All with LOD score limit 30.

Marker positions for each chromosome were constructed using Lep-MAP3 module OrderMarkers2, using only paternally informative markers with recombination1 and interference1 parameters set to 0.000001. The map inflation due to the dependency of adjacent posteriors was reduced by scaling the posteriors by 0.5 in log space (taking the square root) and limiting the minimum posterior probability to 0.1 (scale=0.5 minError=0.1). Maps for each family were examined manually, removing erroneous markers including some distorted markers that passed filtering due to small family sizes.

### Genome scaffolding

Hmel2 scaffolds were manually ordered according to the linkage maps for each of the three *Heliconius melpomene* crosses wherever possible. A small number of misassemblies in Hmel2 were corrected, with scaffolds being split and reoriented where necessary. Not all scaffolds could be ordered based on the linkage maps alone, so Pacific Biosciences reads were also used. PacBio reads were aligned to Hmel2 scaffolds using BWA mem with -x pacbio option [106]. Scaffolds were ordered by manual inspection of spanning reads between scaffolds, identified and summarised by script find_pacbio_scaffold_overlaps.py. Chromosomal positions were assigned by inserting dummy 100 bp gaps between each pair of remaining scaffolds. Although PacBio sequencing could fill gaps between scaffolds, we chose not to do this for these analyses to avoid disrupting Hmel2 linkage map and annotation feature coordinates.

### Recombination rate measurement and permutation testing

CentiMorgan values were calculated using the recombination fraction alone, as the maps were sufficiently fine-scale that mapping functions were not necessary [107]. Per-cross maps (Figure S1) and map statistics (Table 2) were calculated for F1 parents within-species and grandparents for hybrids. Recombination rates were calculated in windows of 1 Mb with 100 kb steps. Chromosomes were tested for reductions in recombination rate in the hybrids compared to *H. melpomene* or *H. cydno* using a bootstrapped Kolmogorov-Smirnov test suitable for discrete data with ties such as recombination counts (ks.boot in the R Matching package [108]), using a one-tailed test for reduced rates in hybrids with 10 000 bootstrap samples, declaring significance at a 0.05 false discovery rate with control for multiple testing (42 tests, with 2 comparisons for each of 21 chromosomes). 1 Mb windows were tested for differences in recombination rate by calculating null distributions of rate differences by permutation of species labels across all offspring, again testing at a 0.05 false discovery rate over 270 000 permutations, controlling for multiple tests with 3 comparisons for each of 2 633 windows. 95% confidence intervals in Figure S2 were calculated by bootstrap, sampling offspring for each species by replacement 10 000 times and calculating cM values, plotting 2.5% and 97.5% quantiles for each window.

### Inversion discovery

PBHoney (in PBSuite version 15.8.24 [109]) was used to call candidate inversions between *H. melpomene* and *H. cydno*, using alignments of PacBio data to ordered Hmel2 scaffolds made with BWA mem with -x pacbio option [106], retaining only primary alignments, and accepting alignments with minimum mapping quality of 30 in Honey.py tails, running on each of four samples (*H. cydno* females, *H. cydno* males, *H. melpomene* females, *H. melpomene* males) separately. Breakpoint candidate sets were compiled together into one file and scaffold positions converted to chromosome positions using script compile_tails.py. PBHoney was run with default options, requiring a minimum of 3 overlapping reads from 3 unique zero-mode waveguides to call a breakpoint candidate. As the *H. cydno* male sample had low coverage, we also ran PBHoney requiring a minimum of 2 reads from 1 zero-mode waveguide and included these tentative candidates where they overlapped with candidates from other samples called with the default settings.

PBHoney was tested for false positives by simulating PacBio reads with pbsim 1.0.3 [110], generating a sample profile using the *H. melpomene* female sample and simulating 15 ‘SMRT cells’ at 5x coverage each. Simulated data was then aligned with BWA and inversions called with PBHoney as above.

Trio assemblies were aligned to Hmel2_ordered using NUCmer from the MUMmer suite ([111]; version 3.23), followed by show-coords with -Tlcd options, to produce tab-separated output including scaffold lengths, percentage identities and directions of hits.

Script detect_inversion_gaps.py was used to integrate the PBHoney inversion candidates with the linkage maps, trio alignments and *H. melpomene* annotation (from Hmel2). As this data is being used to rule out inversions in regions without recombinations, PBHoney inversion candidates were rejected if at least one recombination for the same species as the candidate was contained within the inversion. PBHoney candidates were also rejected if there was a trio scaffold alignment spanning the candidate inversion, with spanning defined as extending more than half the length of the candidate inversion in either direction. Finally, candidates shorter than 1 kb were rejected, as linkage disequilibrium between SNPs separated by 1 kb or less in *H. melpomene* rises above background levels [94] and so inversions are unlikely to be required to maintain linkage. The retained inversion candidates were then combined into groups by overlap.

Each group of overlapping inversion candidates was then classified as follows (Figure 3, Table 5, Figures S8-14): *Split reads and trio assembly*, group has at least one PBHoney inversion candidate and at least one trio scaffold with forward and reverse alignments either side of an inversion breakpoint; *Split reads only,* group has at least one PBHoney inversion candidate in at least one sex, but no matching inverted trio scaffolds; *Split reads in one species*, *trio assembly in both*, group has at least one PBHoney inversion candidate in at least one sex of only one species, but trio assembly has inverted scaffolds in at least one sex in both species. These classifications do not cover whether there are multiple contigs across the candidate inversion (see Table 5, Figures S8-14) or whether trio scaffolds with alignments that span whole PBHoney inversion candidates or cross candidate breakpoints (see Figures S8-14).

### Population genetics statistics

*F*_*ST*_ and *f*_*d*_ were calculated within and around candidate inversions following Martin et al. 2013 [84] and Martin et al. 2015 [95], using all 4 *H. melpomene rosina*, 4 *H. cydno chioneus*, 4 *H. melpomene* French Guiana and 2 *H. pardalinus* samples from Martin et al. 2013 [84]. Samples were aligned to Hmel2 using bwa mem v0.7.12 using default parameters and genotypes were called GATK v3.4 HaplotypeCaller using default parameters except for setting heterozygosity to 0.02. For each candidate inversion, 11 windows equal to the size of the inversion were generated, one for the inversion itself and five either side of the inversion, except where candidates were at the ends of chromosomes. Statistics were calculated for each window with scripts popgenWindows.py and ABBABABAwindows.py in GitHub repository genomics_general (https://github.com/simonhmartin/genomics_general).

### H. erato analysis

The *H. erato* version 1 genome assembly was downloaded from LepBase (http://ensembl.lepbase.org/Heliconius_erato_v1) and aligned to the ordered Hmel2 scaffolds with LAST version 744 [112]. Scaffolds and linkage maps were compared with bespoke scripts Hmel2_Herato_maf.py, compile_Herato_maps.py and Hmel2_Herato_dotplot.R.

## Acknowledgements

Thanks to Owen McMillan, Adriana Tapia, Liz Evans, Maria-Clara Melo, Janet Scott, Bas van Schooten, Adrea Gonzalez-Karlsson, Moises Abanto, Tim Thurman and all staff at the Smithsonian Tropical Research Institute for support and assistance with insectary work. RAD protocol developed by Paul Etter at the University of Oregon and further enhanced by Pablo Fuentes-Utrilla and Cathlene Eland at Edinburgh Genomics. PacBio sequencing was carried out by Karen Oliver at the Sanger Institute. Thanks to Jenny Barna for computing support. Analyses were performed using the Darwin Supercomputer of the University of Cambridge High Performance Computing Service (http://www.hpc.cam.ac.uk/), provided by Dell Inc.

## Supporting Information Legends

**Figure S1. Marey maps of recombinations for each cross separately.** Crosses listed in Table 2, Table S1 and Table S2. *H. melpomene*, red; *H. cydno*, blue; *H. cydno* x *H. melpomene* hybrids, green. Chromosomes 1-21 of *H. melpomene* genome assembly version 2 (Hmel2) shown against cumulative cM values for each set of crosses.

**Figure S2. Recombination rates in cM/Mb for each Hmel2 chromosome.** Rates in 1 Mb windows sliding by 100 kb. *H. melpomene*, red; *H. cydno*, blue; hybrids, green. Lines show true values; shaded areas show 95% bootstrap confidence intervals, 10 000 iterations.

**Figure S3. Gaps in linkage maps where lack of precise recombination data cannot rule out presence of inversions.** Each page shows data for one chromosome with *H. cydno* in blue and *H. melpomene* in red. Heights and lengths of lines show gap lengths.

**Figure S4. Histograms of gap lengths for** ***H. cydno*** **and** ***H. melpomene***,.

**Figure S5. Probability of detecting random inversions of sizes from 10 kb to 1.5 Mb given existing linkage maps for** ***H. melpomene*** **and** ***H. cydno***.

**Figure S6. Histograms of raw read lengths for Pacific Biosciences sequencing.** *H. cydno* females, blue; *H. cydno* males, light blue; *H. melpomene* females, red; *H. melpomene* males, orange.

**Figure S7. Histograms of base depths across the genome after alignment of raw PacBio reads to** ***H. melpomene*** **genome assembly Hmel2.** Colours as for Figure S6.

**Figures S8-14. Full evidence for each candidate inversion group, separated into the classes shown in Figure 3 and Table 5. S8**, *H. cydno*, Split reads and trio assembly. **S9**, *H. cydno*, Split reads only. **S10**, *H. melpomene*, Split reads and trio assembly. **S11**, *H. melpomene*, Split reads only. **S12**, Both species, Split reads and trio assembly. **S13**, Both species, Split reads only. **S14**, Both species, Split reads in one species, trio assembly in both.

Each page shows linkage map, split read, trio assembly and population genetics evidence for each candidate inversion group, across a region of the Hmel2 genome assembly. Black lozenge, candidate inverted region (white text in lozenge shows length of region). **Genome**, contigs from Hmel1 and Hmel2 shown in alternating dark and light grey (labels show Hmel1 scaffold names and Hmel2 contig names). Thin black lines in Hmel2 contigs show repeat-masked regions of the genome. Red bars show features from the Hmel2 annotation. **Linkage map**, SNPs from linkage maps shown as circles connected by Marey map lines. Thick lines show regions with no recombination; thin lines span regions between SNPs with different markers, indicating a recombination happens somewhere along the line but the position cannot be resolved any further. *H. cydno*, blue; *H. melpomene*, red; hybrids, green. **PBHoney Candidates**, rounded lines show candidate inversions called by PBHoney; number to the right shows number of reads supporting each candidate. Light shaded rectangles show ranges of these inversions across the plot for comparison with trio alignments and contig boundaries. *H. cydno* females, blue; *H. cydno* males, light blue; *H. melpomene* females, red; *H. melpomene* males, orange. **Trio** strips, alignments of trio scaffolds to candidate inversion regions. Arrowed lines show individual alignments; shaded rectangles enclosing arrowed lines show all alignments for a particular scaffold, with scaffold name to the right of each shaded rectangle. Colours as per PBHoney candidates. **Trio Inverted**, trio scaffolds with multiple alignments across the candidate inversion, with forward and reverse alignments either side of candidate breakpoints. **Trio Spanning**, trio scaffolds with single alignments spanning any one PBHoney candidate. **Trio Edges**, trio scaffolds with alignments spanning one candidate breakpoint, but not the whole inversion. **Population genetics**, *F*_*ST*_ (magenta) and *f*_*d*_ (green) for windows across the candidate inversion. Black lozenge, inverted region, matching lozenge in main plot. Length and number label of grey lozenges at the bottom show number of sites used for calculations in each window. Open circles show windows were calculation is not possible (either no sites present, or D<0 for *f*_*d*_).

**Figure S15. Oxford grids for ordered Hmel2 scaffolds and** ***H. erato*** **scaffolds.** Collinear regions in red; inverted regions in blue. Scaffold boundaries separated by grey lines. Black and dark grey lines on right and lower axes show alternating markers from linkage maps; light grey shaded rectangles show these markers across the plot.

**Table S1. Full sample details for** ***H. melpomene*** **and** ***H. cydno*** **crosses.** All samples were RAD sequenced except for underlined samples, which were whole genome sequenced and trio assembled. Samples shaded in grey were rejected due to low coverage or many erroneous genotypes. ENA = European Nucleotide Archive, http://www.ebi.ac.uk/ena.

**Table S2. Full sample details for** ***H. cydno*** **x** ***H. melpomene*** **hybrid crosses.** Each cross involved multiple backcrosses. Many parents were not preserved or sequenced. Samples shaded in grey were rejected due to low coverage or many erroneous genotypes. ENA = European Nucleotide Archive, http://www.ebi.ac.uk/ena.

**Table S3. Counts of raw and filtered PBHoney candidate inversions.**

